# An electrochemical approach for rapid, sensitive, and selective detection of dynorphin

**DOI:** 10.1101/2023.02.01.526701

**Authors:** Sineadh M. Conway, Chao-Cheng Kuo, Woodrow Gardiner, Rui-Ni Wu, Loc V. Thang, Graydon B. Gereau, John R. Cirrito, Carla M. Yuede, Jordan G. McCall, Ream Al-Hasani

**Affiliations:** Center of Clinical Pharmacology, University of Health Sciences and Pharmacy and Washington University in St. Louis, USA St. Louis, MO USA; Department of Anaesthesiology Washington University in St. Louis, USA; Washington University Pain Center, Washington University in St. Louis, MO, USA; Department of Neurology, Washington University in St. Louis, USA; Department of Psychiatry, Washington University in St. Louis, USA

**Keywords:** electrochemistry, voltammetry, opioids, dynorphin

## Abstract

The endogenous opioid peptide systems are critical for analgesia, reward processing, and affect, but research on their release dynamics and function has been challenging. Here, we have developed microimmunoelectrodes (MIEs) for the electrochemical detection of opioid peptides using square-wave voltammetry. Briefly, a voltage is applied to the electrode to cause oxidation of the tyrosine residue on the opioid peptide of interest, which is detected as current. To provide selectivity to these voltammetric measurements, the carbon fiber surface of the MIE is coated with an antiserum selective to the opioid peptide of interest. To test the sensitivity of the MIEs, electrodes are immersed in solutions containing different concentrations of opioid peptides, and peak oxidative current is measured. We show that dynorphin antiserum-coated electrodes are sensitive to increasing concentrations of dynorphin in the attomolar range. To confirm selectivity, we also measured the oxidative current from exposure to tyrosine and other opioid peptides in solution. Our data show that dynorphin antiserum-coated MIEs are sensitive and selective for dynorphin with little to no oxidative current observed in met-enkephalin and tyrosine solutions. Additionally, we demonstrate the utility of these MIEs in an *in vitro* brain slice preparation using bath application of dynorphin as well as optogenetic activation of dynorphin release. Future work aims to use MIEs *in vivo* for real-time, rapid detection of endogenous opioid peptide release in awake, behaving animals.

## Introduction

Endogenous opioid peptides including endorphins, enkephalins, dynorphins, and nociceptin, are critical for analgesia, reward processing, and affect. Research on their function in the modulation of these behaviors has been challenging for several reasons. Opioid peptides have similar amino acid sequences making them difficult to distinguish from one another. For example, dynorphin-related opioid peptides share a common N-terminal sequence with leu-enkephalin, and C-terminal residues are conserved with dynorphin A 1-17. Furthermore, opioid peptides are rapidly cleaved by peptidases, undergo posttranslational modifications, and are often found at concentrations several orders of magnitude lower when compared to classical neurotransmitters.^1,2^ These properties require detection and quantification techniques with a high spatial and temporal resolution, in conjunction with high sensitivity and selectivity. Together these requirements have made it difficult to reliably and consistently detect dynamic changes in opioid peptides.

Several useful techniques currently exist for the detection and quantification of opioid peptides, but each carries unique limitations. Techniques using radioimmunoassays or enzyme-linked immunosorbent assays (ELISA) rely on the selectivity of antibodies to distinguish and quantify different opioid peptides and often require larger sample volumes resulting in low temporal resolution. Microdialysis has also been paired with liquid chromatography/mass spectrometry, which has improved peptide recovery and detection as well as increased the temporal resolution to tens of minutes and does not rely on antibody-based results ^3 4 5 6 7 8^. While these advances have greatly improved quantification accuracy, there is still a need for improved temporal resolution.

A promising advance in the detection and quantification of opioid peptides employs the electrochemical technique of voltammetry, which has traditionally been used for the detection of monoamines like dopamine.^9–15^ This pioneering work uses a multiple-scan-rate voltammetry waveform for real-time detection of met-enkephalin in adrenal tissue and in the dorsal striatum of anesthetized and freely moving rats.^16,17^ Met-enkephalin consists of electroactive tyrosine and methionine residues that oxidize at unique electrical potentials and are detected as an increase in current. This electrochemical approach advanced opioid peptide detection in several arenas. First, the small carbon fiber microelectrode is acutely implanted into the brain which creates less gliosis than traditional microdialysis probes and can detect opioid peptide release much closer to possible release sites.^16,17^ Secondly, and arguably most importantly, voltammetry vastly decreases the sampling rate to a subsecond temporal resolution which is particularly important in quantifying opioid peptides that are rapidly degraded or cleaved after release. Despite these important advances, this voltammetric technique is limited to monitoring amino acid components of opioid peptides, making it difficult to selectively distinguish opioid peptides such as dynorphins that do not have a unique electroactive residue.

Building on this work, we employ a voltammetric approach using microimmunoelectrodes (MIEs) to monitor dynamic changes in dynorphin concentrations, with the goal of developing a rapid, sensitive, and selective technique for the detection and quantification of dynorphin. MIEs have previously been successfully used to quantify β-amyloid in mouse models of Alzheimer’s disease.^18^ Here, we use square-wave voltammetry to oxidize the tyrosine residue found in dynorphin, which is detected as an increase in current. Importantly, tyrosine is a common amino acid in many neurotransmitters and neuropeptides including opioid peptides and so to add selectivity for our peptide of interest, the carbon fiber microelectrode is first coated with antiserum selective for dynorphin. Thus, an increase in current is a result of the oxidation of tyrosine residues in dynorphin. Overall, our work shows MIEs can be used for selective and sensitive monitoring of dynorphin concentrations at the sub-femtomolar to picomolar range on the timescale of seconds. We validate its use *in vitro* for the monitoring of pharmacological- and photostimulation-induced dynorphin release in mouse brain slice preparations.

## Results and Discussion

The precise role of endogenous opioid peptides in behavior has been difficult to discern because of a lack of tools to measure dynamic concentration changes in a sensitive and selective manner. Here, we have developed a voltammetric technique for the rapid measurement of dynorphin release dynamics in a sensitive and selective manner. This approach can be widely implemented and provides a previously unachievable spatiotemporal resolution for directly measuring dynorphin release.

Voltammetry is a common electrochemical technique used to measure electroactive species. Dynorphin peptides can be measured by using the intrinsic electroactivity of the N-terminal tyrosine residue. Tyrosine consists of a phenolic group that can be oxidized at the surface of a microelectrode. When oxidation occurs, the electroactive species, tyrosine, releases an electron that is detected as current and is directly proportional to the concentration present. In this case, tyrosine oxidizes at ∼0.65 V. To selectively measure tyrosine from dynorphin, the electrode must first be coated with dynorphin antiserum. Figure 1A shows the timeline of MIE preparation across days. After electrode fabrication, the exposed carbon fiber (Figure 1B) is electrically and chemically activated to allow for antibody attachment. The electrode is then incubated in the antiserum overnight. On day 2, the attachment step is completed, and the electrode is incubated in bovine serum albumin (BSA) to block any non-specific binding sites. At this point, the MIE is ready for calibration and experimental recordings. Figure 1C shows the process by which dynorphin is measured at the electrode surface. First, a square wave electrical potential (Figure 1D) is applied to the MIE from 0 to 1 V. If dynorphin is present at the electrode surface, its tyrosine residue will oxidize at ∼0.65V, transferring an electrode to the MIE that is detected as current. This increase in current is referred to as peak oxidative current and is directly proportional to the concentration of tyrosine present (Figure 1E).

**Figure 1.**
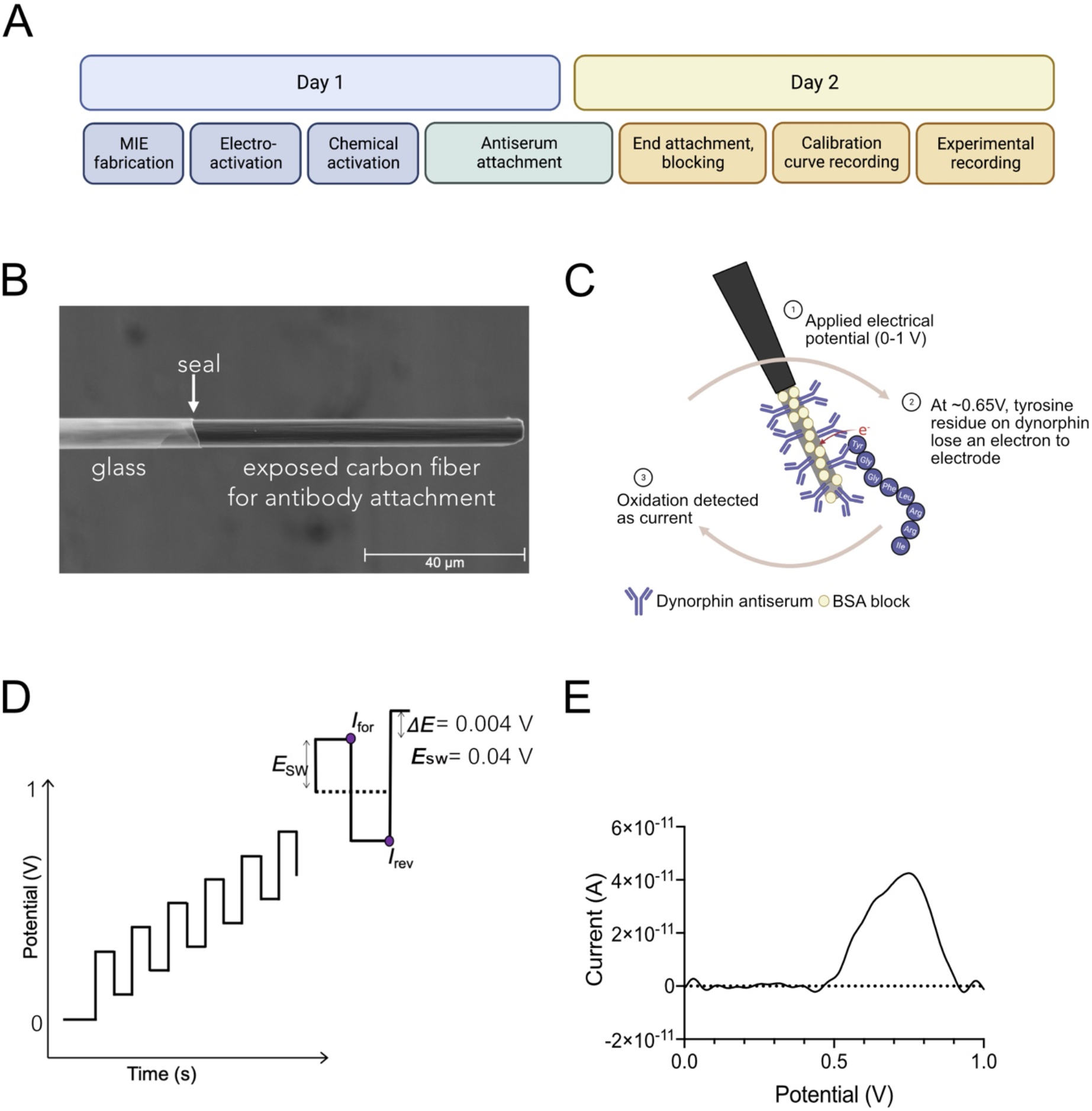
Methodology for MIE preparation and voltammetry recording. A. Timeline for MIE fabrication and recording. B. Scanning electron microscopy image of MIE tip with exposed carbon fiber for antiserum attachment. C. Conceptual representation of MIE voltammetric approach. D. Square-wave voltammetry signal used to apply an electrical potential to the MIE electrode surface. (E_SW_=0.04V, ΔE=0.004V). Current (I) is measured twice during each step (forward and reverse). E. Example of processed current data from individual MIE exposed to dynorphin.

To confirm that the peak oxidative current measured is from tyrosine residues on dynorphin, each MIE is calibrated against different concentrations of dynorphin, tyrosine, and metenkephalin (an opioid peptide with a different amino acid sequence but still contains a tyrosine residue). In this manner, we are testing both the selectivity and sensitivity of our dynorphin MIE. Figures 2A-C show representative voltammograms of a single electrode calibrated to dynorphin, met-enkephalin, and tyrosine, respectively. Importantly, peak oxidative current increases with increasing concentrations of dynorphin, and the peak currents observed from exposing the MIE to met-enkephalin and tyrosine are extremely low. Thus, we can conclude that this MIE is sensitive and selective for dynorphin. Figure 2D shows the calibration of 10 dynorphin MIEs against dynorphin, met-enkephalin, and tyrosine across a range of concentrations. A mixed-effect model of analysis revealed a main effect of concentration, a main effect of analyte, and a significant interaction between concentration and analyte. Overall, the peak oxidative current from dynorphin was significantly different from met-enkephalin and tyrosine at 0.0005, 0.001, 0.5, and 1 pM. Importantly, the MIEs show an increase in peak oxidative current only across increasing concentrations of dynorphin, which confirms the MIEs sensitivity to dynorphin. A small current is still measured when tested against met-enkephalin and tyrosine, but it does not increase with increasing concentration. Thus, the MIEs are not sensitive to tyrosine alone or met-enkephalin and are most selective for dynorphin. This calibration step is necessary to test the efficiency of each individual electrode prior to use in additional experiments. It is not uncommon to have poorly performing electrodes that do not calibrate to dynorphin, and these MIEs are discarded. Poor performance can occur for a multitude of reasons, such as a crack in the glass, an insufficient seal between the glass and the exposed carbon fiber, carbon fiber length, or inadequate antiserum binding. Additionally, each MIE is unique in the exact length and efficiency of antiserum attachment. Thus, there are individual differences in peak oxidation current measured across individual electrodes for similar concentrations of each analyte. Therefore, individual calibration is necessary prior to use to validate the efficacy of the MIE and to estimate current-to-concentration conversions.

**Figure 2.**
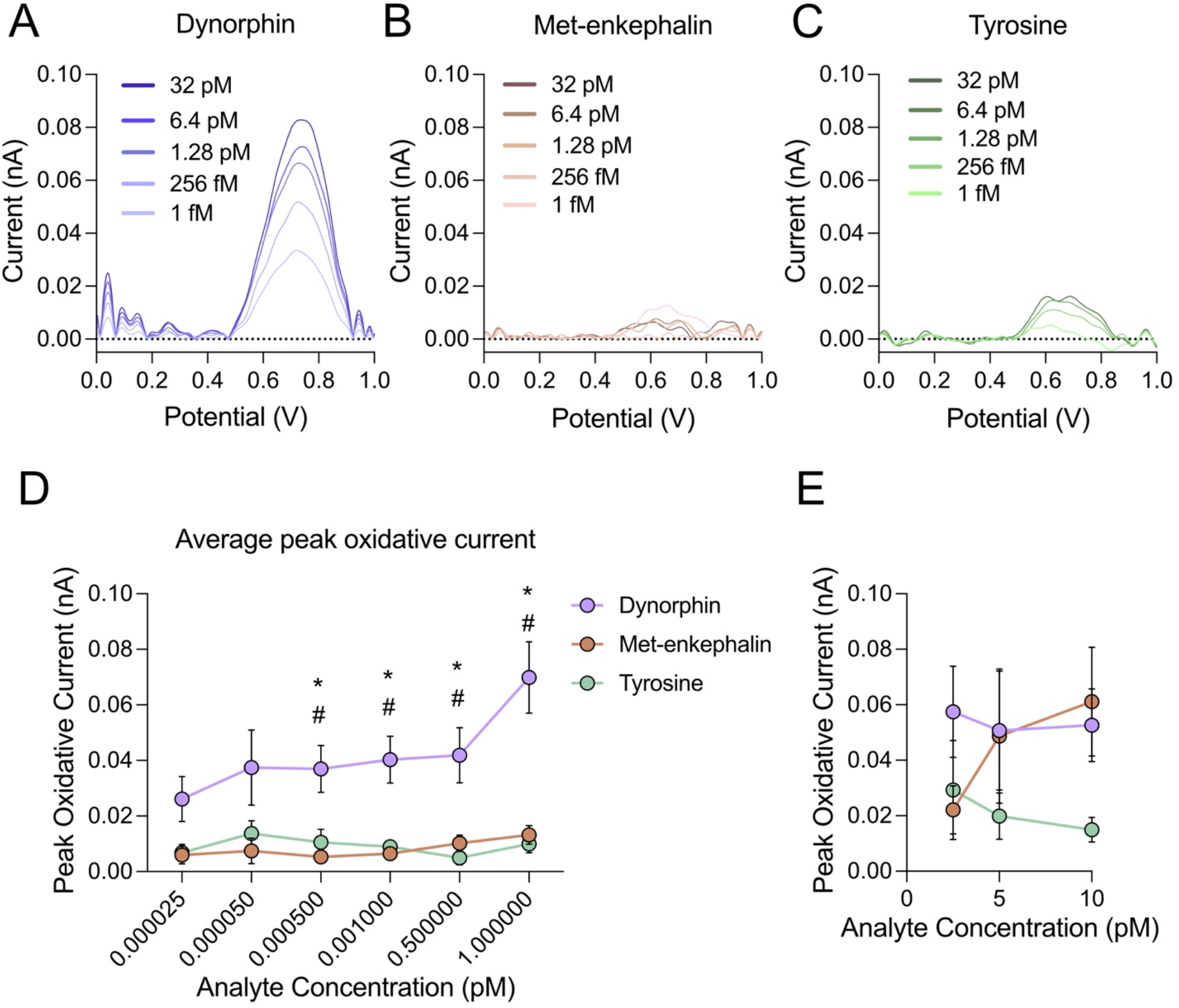
Validation of dynorphin MIE sensitivity and selectivity. A. Representative MIE calibration to dynorphin in solution with peak oxidative current increasing with increased dynorphin concentration, indicative of sensitivity. B. The same electrode depicted in A does not show sensitivity or selectivity to met-enkephalin. C. The same electrode depicted in A and B does not show sensitivity or selectivity to tyrosine. D. Average peak oxidative current from 10 MIE electrodes tested against dynorphin, met-enkephalin, and tyrosine is sensitive and selective for dynorphin in the sub-femtomolar to low picomolar range. The same MIE electrodes depicted in D are not selective for dynorphin at higher picomolar concentrations. \**p*<0.05 dynorphin vs. met-enkephalin. *#p*<0.05 dynorphin vs. tyrosine.

Importantly, the increase in peak oxidative current across increasing concentrations of dynorphin is observed at the sub-femtomolar to picomolar range (Figure 2D) but not at higher concentrations (Figure 2E). This is ideal as we predict endogenous concentrations of dynorphin to be within this range, as previous work has shown that evoked dynorphin release is in the picomolar range,^19^ so it is likely that the non-evoked, endogenous concentration range will be lower. Furthermore, these previous measurements were collected from multiple slices in larger volumes of artificial cerebral spinal fluid (aCSF) using ELISA quantification, whereas the measurements occurring at the MIE surface will be in a much smaller area. Overall, these data show that MIEs are an ideal tool for measuring low concentrations of dynorphin.

Following confirmation of MIE selectivity and sensitivity to dynorphin, individual MIEs were tested using the same electrical setup we use for acute brain slice electrophysiology for validation of future *in vitro* recordings. Figure 3A shows the experimental timeline for the MIE recordings. First, priming scans are performed in which the square wave is applied for 30 minutes to habituate the MIE to the experimental environment. This is similar to other voltammetry experiments that need priming scans to equilibrate the current and reduce drift. After 30 minutes of priming scans, current was measured following square-wave application of electrical potential (0 to 1 V, 30 Hz) every 15 seconds during bath application of aCSF solution and 1 pM dynorphin in aCSF. Prior to experimental recording, individual MIEs are calibrated to confirm sensitive and selective dynorphin detection (Figure 3B). Figure 3C shows an individual MIE measurement of peak oxidative current measured across each square-wave voltammetry scan. Peak oxidative current is then binned into 5-minute time blocks (Figure 3D) to compare measurements during bath application of aCSF alone (gray circles) to bath application of 1 pM dynorphin in aCSF (purple circles). A repeated measures one-way analysis of variance (AVONA) revealed a significant difference between peak oxidative current during dynorphin application and aCSF alone (p<0.001). Figure 3E shows the average calibration of two MIEs for bath application recordings. Average peak oxidative current across two MIEs to bath application of aCSF alone and 1 pM dynorphin in aCSF is shown in Figure 3F. To compare peak oxidative current across the recording session, measurements are binned into 5 min time blocks (Figure 3G). Repeated measures one-way ANOVA revealed a significant increase in average peak oxidative current during 1 pM dynorphin + aCSF bath application (time blocks 2-4) compared to aCSF application alone (time blocks 1, 5, and 6; p<0.0001). Importantly, the peak oxidative current measurements are dynamic with increased signal observed during dynorphin application that decreases following washout. These data highlight the ability to use MIEs *in vitro* and confirm the dynamic signal.

**Figure 3.**
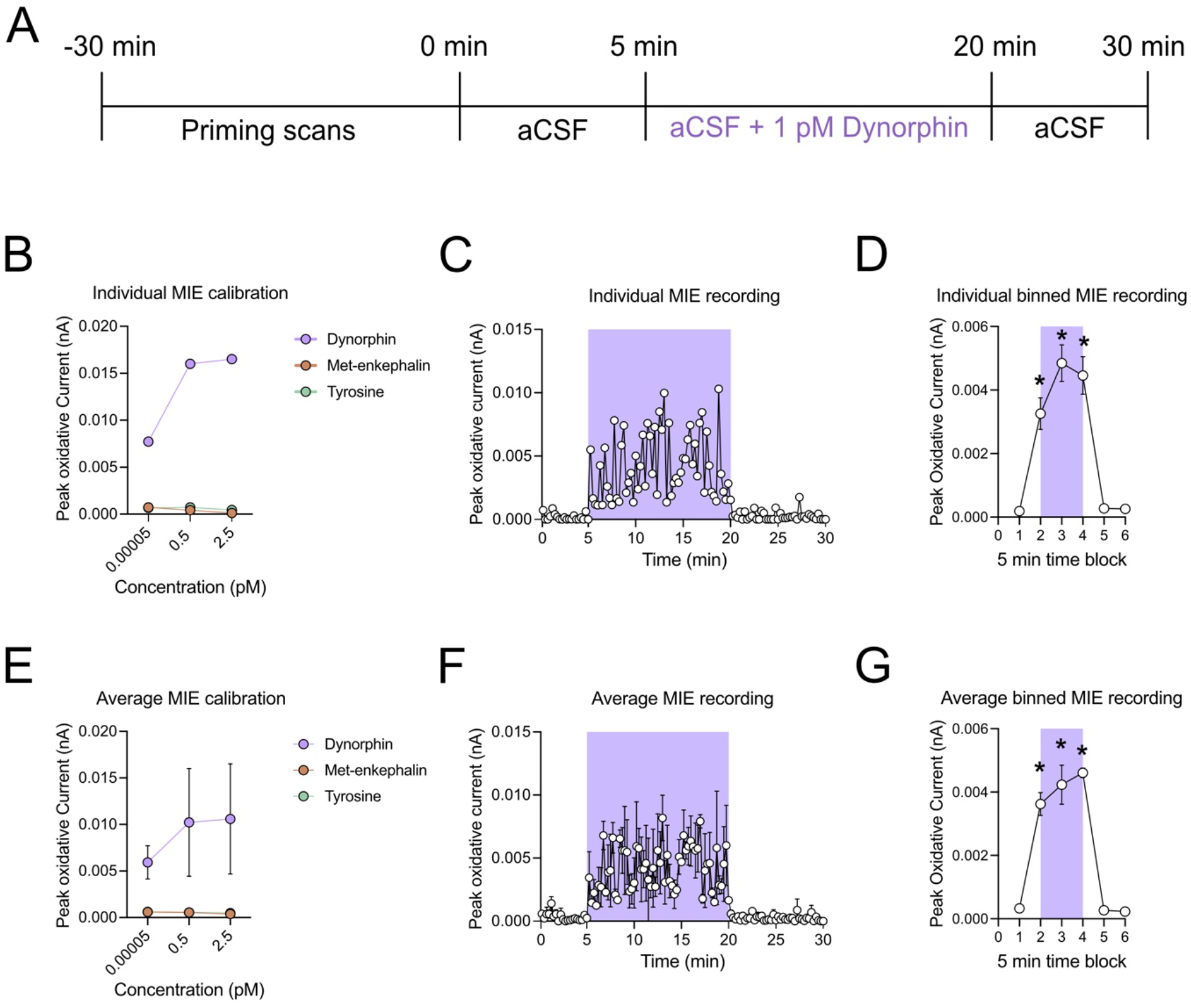
*In vitro* validation of dynorphin MIE detection. A. Experimental timeline for bath application of dynorphin. Following 30 minutes of priming scans to habituate the electrode, aCSF was bath applied for 5 minutes followed by bath application of 1 pM dynorphin in aCSF for 15 minutes and aCSF for 10 minutes. B. Individual example of MIE calibration to recording to confirm sensitive and selective detection of dynorphin, observed as an increase in peak oxidative current to increasing concentrations of dynorphin only. C. Individual example of peak oxidative current measured from MIE across recording session. D. Individual MIE recording measurement of peak oxidative current binned by 5 min. E. Average calibration of two MIEs used in bath application experiment. F. Average peak oxidative current measured from two MIEs across recording session. G. Averaged MIE recording measurement of peak oxidative current binned by 5 min. \**p*<0.0001 compared to aCSF bins (5 min time blocks 1, 5, and 6). Purple shading indicates bath application of dynorphin.

Next, we determined whether MIEs can detect optically-evoked dynorphin release. To do so, we injected either AAV5-DIO-ChR2-eYFP or AAV5-DIO-eYFP into the NAc shell (from Bregma: anteroposterior 1.3 mm, mediolateral 0.5 mm, dorsoventral -4.2 mm) of preprodynorphin-cre (pDyn-Cre) mice (Figure 4A). After allowing 30 days for the virus to express (Figure 4B), acute brain slices were prepared, and the MIE was positioned in the medial NAc shell (Figure 4C). Following 30 min of priming scans, current was measured following repetitive square-wave application of 0 to 1 V every 15 seconds before, during, and after photostimulation (10 Hz, 10 ms pulse width) (Figure 4D). Prior to the experimental recording, individual electrodes are calibrated (Figure 4E). Figure 4F shows an increase in peak oxidative current during optogenetic stimulation. A repeated measures one-way analysis of variance (ANOVA) of the average peak oxidative current before, during, and after photostimulation (Figure 4G) revealed a significant increase in peak oxidative current during the photostimulation period (time blocks 2-4; p<0.001) compared to before (time block 1) and after (time blocks 5 and 6).

**Figure 4.**
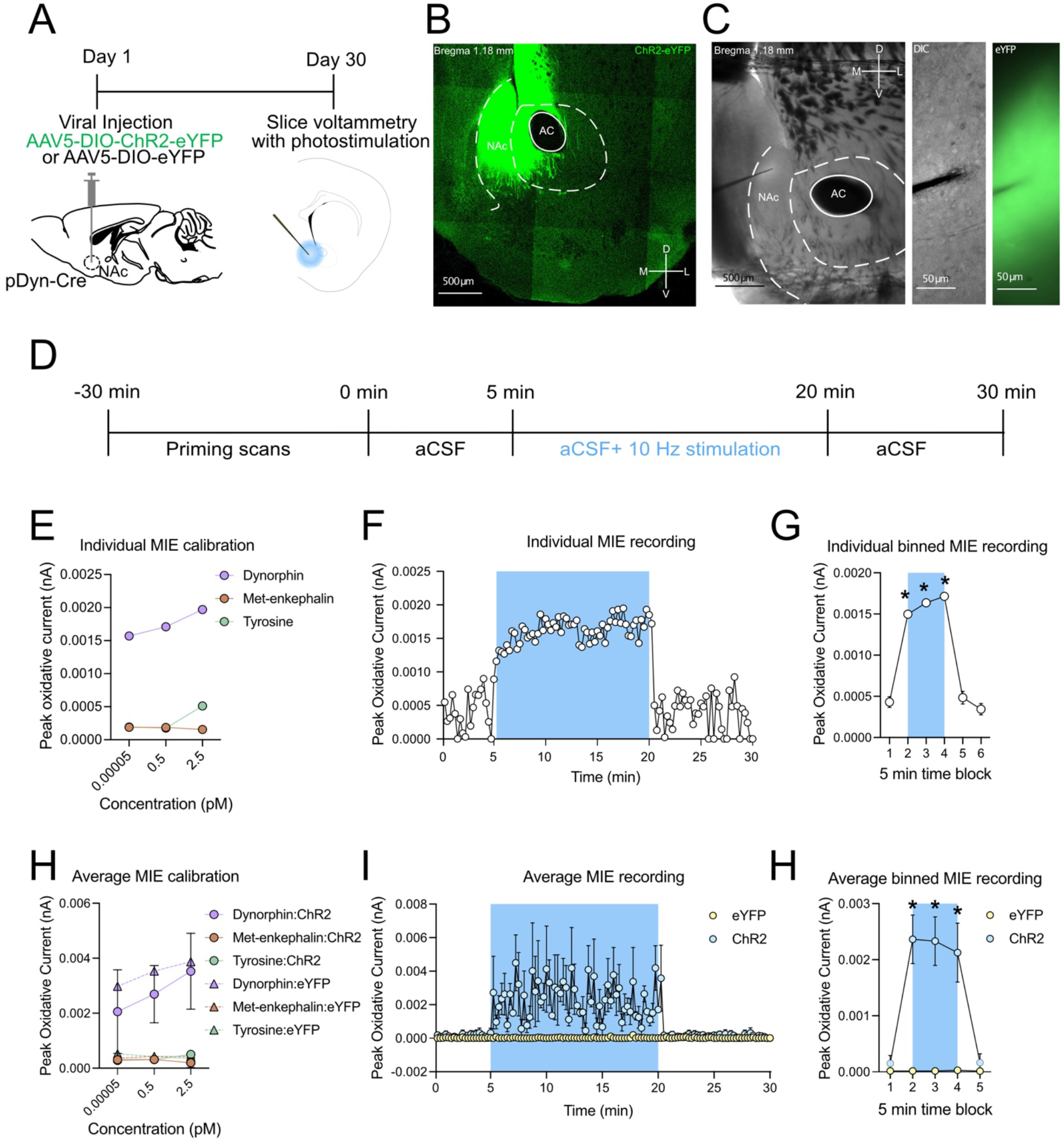
*In vitro* detection of optogenetically evoked dynorphin release in the nucleus accumbens. A. Experimental timeline for expression of channelrhodopsin or eYFP in the NAc of pDyn-Cre mice. B. Representative image of channelrhodopsin expression in the NAc. C. Image of MIE in brain slice. D. MIE recording timeline. E. Individual example of MIE calibration prior to recording to confirm sensitive and selective detection of dynorphin, observed as an increase in peak oxidative current to increasing concentrations of dynorphin only. F. Peak oxidative current measured from individual MIE across recording session. G. Individual MIE recording measurement of peak oxidative current binned by 5 min. H. Average MIE calibrations for three MIEs used in recordings from ChR2 mice and two MIEs using in recordings from eYFP mice. I. Average peak oxidative current measured during ChR2 and eYFP recording sessions. J. Average peak oxidative current from ChR2 and eYFP recording sessions binned by 5 min. \**p*<0.003. Blue shading indicates 10 Hz photostimulation. NAc=nucleus accumbens; AC=anterior commissure.

To confirm the photostimulation-induced increase in peak oxidative current in pDyn-Cre brain slices, the same experiment was performed in pDyn-Cre mice transfected with eYFP. Figure 4H shows the average calibration results from the three MIEs used in ChR2 brain slice recordings and two MIEs used in eYFP brain slice recordings. Overall, all electrodes were sensitive and selective for dynorphin as shown by an increase in peak oxidative current to increasing concentrations of dynorphin (purple circles and triangles). The average time course measurements (Figure 4I) show that peak oxidative current increases during 10 Hz photostimulation in the ChR2 brain slices. The ChR2 group shows a significant increase in peak oxidative current compared to eYFP controls during the photostimulation period, as revealed by a repeated measures two-way ANOVA (Figure 4H, p<0.003). In sum, these data confirm that MIEs can detect optically evoked dynorphin release in the medial NAc shell. Importantly, the signal is dynamic, is not corrupted by photovoltaic artifacts, and peak oxidative current returns to baseline levels once the photostimulation is removed.

Overall, the use of MIEs for dynorphin detection provides a previously unachievable measurement of endogenous dynorphin in brain slices. Detection of neuropeptides in acute brain slices is notoriously difficult, the sensitivity of this new approach provides much needed access to observing endogenous opioid peptide release. Remarkably, our data suggests that acute NAc slices maintain the ability to release dynorphin for as long as 15 minutes, although it is possible that this duration is prolonged due to a washout of endogenous peptidases.

Importantly, this technique provides a high spatial and temporal resolution for measuring low concentrations of dynorphin. The equipment is relatively inexpensive, enabling its cost-effective implementation, particularly for labs that have existing voltammetry or electrophysiology equipment that can be adapted for these purposes. The higher costs in applying this technique come from the time required to prepare the MIE electrodes. Each MIE must be fabricated and calibrated prior to use in experimental recordings. MIEs are extremely fragile so careful handling during the entire process is vital. Furthermore, MIEs can only be used acutely. Thus, MIEs are ideal for experiments with shorter timescales of 1-2 hours. This MIE technology provides an excellent means to measure endogenous release of dynorphin, and future studies will validate its use for *in vivo* measurements of dynorphin release. Importantly, this technique can be applied to other opioid peptides and neuropeptides that contain electrochemically active amino acid residues, thus providing a previously unachieved tool for direct electrochemical measurements of peptide release.

## Methods

### Materials

Carbon fibers (5um in diameter) were obtained from GoodFellow Corp. (Boulder City, NV). Single-barrel borosilicate capillary glass was obtained from A-M Systems (Sequim, WA). 1-ethyl-3-[3-dimethylaminopropyl] carbodiimide hydrochloride (EDC) was obtained from Life Technologies (Carlsbad, CA) and N-hydroxysulfosuccinimide (NHSS) was obtained from Bioworld (Dublin, OH). Ethanolamine was purchased from Chem-Impex (Wood Dale, IL). Dynorphin antiserum was purchased from US Biological (Salem, MA). Dynorphin 1-8 peptide was purchased from New England Peptide (Gardner, MA). Met-enkephalin acetate salt hydrate and bovine serum albumin solution were purchased from Sigma-Aldrich (St. Louis, MO). Tyrosine was purchased from TCI Chemicals (Tokyo, Japan). The image of the MIE (Figure 1B) was taken on a Thermofisher Quattro S Environmental Scanning Electron Microscope (ESEM).

### Microimmunoelectrode fabrication

MIEs were prepared as previously described^18^. A single carbon fiber was aspirated into a glass capillary tube (0.4 mm inner diameter, 10 in length) using a vacuum. Each filled capillary tube was pulled into two microelectrodes using a pipette puller (Narishige PE-22). The carbon fiber was attached to an insulated silver wire using conductive silver adhesive paste (Ted Pella, Inc.), and the connection between the wire and glass was sealed using heat shrink tubing (Altex). The exposed carbon fiber was cut with a scalpel to a length of 30-100 um. Microelectrodes were pretreated using a triangular waveform from 0 to 3 V for 70 s in PBS to enhance antibody binding. The carboxylic groups on the carbon fiber surface were electro-activated by incubation in 0.4 M 1-ethyl-3-[3-dimethylaminopropyl] carbodiimide hydrochloride (EDC; Life Technologies) and 0.1 M and N-hydroxysulfosuccinimide (NHSS; Bioworld) solutions for 1 hr to create a semistable reactive amine NHS ester. The electo-activated microelectrodes are then incubated in dynorphin antiserum at 4°C overnight. Following antiserum attachment, MIEs are placed in 0.01% ethanolamine to deactivate reactive amine sites. Lastly, MIEs are incubated in 0.1% BSA solution for 1 hr to block any nonspecific protein-binding sites.

### MIE calibration testing

Calibration of the MIEs was performed using a BASI potentiostat and software. All calibration testing was done using a conventional 3-electrode cell comprised of the working MIE electrode, an Ag/AgCl reference electrode (3M NaCl), and a platinum wire counter electrode. First, a square wave is applied to the MIE 50 times to habituate the MIE (priming scans). An electrical potential was applied in a stepwise function from 0 to 1 V at a frequency of 30 Hz (Increase E = 0.004 V; Amplitude = 0.04 V). Next, dynorphin, met-enkephalin, and tyrosine solutions were prepared in concentrations ranging from sub-femtomolar to picomolar in PBS. Each electrode was exposed to all solutions at random, and square-wave voltammetry was used to monitor the response of the electrode. The same square wave parameters used to prime the MIE was applied 5 times for each individual sample with 5 s rest time between each scan. The MIE response was calculated by averaging the peak oxidative current from all 5 potential sweeps.

### Square-wave voltammetry analysis

The current output data obtained were analyzed using a custom R-Studio script. First, the raw current data is smoothed using Fourier transformation. Second, the second-order derivative of the smoothed data is used to measure the peak oxidative current between 0.6 and 0.8 V. Peak oxidative current is measured for each voltage scan.

### In vitro validation experiment

To test the use of MIEs on the slice rig (Figure 3), an MIE was positioned in the slice bath without a brain slice present. aCSF was bath applied throughout the entire recording session. A square wave was applied from 0 to 1 V at a frequency of 30 Hz (Incre. E = 0.004 V; Amplitude = 0.04 V) every 15 seconds for 30 min to prime, or habituate, the MIE to the brain slice environment. Next, the same square wave was applied for an additional 5 min and peak oxidative current was measured from each electrical potential scan as the baseline. Immediately following baseline, 1 pM dynorphin in aCSF was bath applied to the MIE was 15 minutes with simultaneous square wave application. Next, aCSF alone was applied for 10 minutes with continued square wave application. Overall, each individual square wave lasted ∼8sec with 5 seconds in between each scan.

### Acute brain slice experiment

Preprodynorphin-IRES-Cre mice (pDyn-Cre) mice were used. Mice were unilaterally injected with 250nL of AAV5-hSyn-DIO-ChR2-eYFP or AAV5-hSyn-DIO-eYFP in the NAc. Mice were kept in their homecage for 4 weeks to allow for recovery and viral expression prior to testing. Animals were deeply anesthetized via intraperitoneal injections of anesthetic mixture containing ketamine, xylazine and acepromazine, then were perfused with ice-cold slicing artificial cerebrospinal fluid (aCSF) that consisted of the following (in mM): 93 N-methyl-D-glucose (NMDG), 2.5 KCl, 1.25 NaH_2_PO_4_, 0.5 CaCl_2_, 20 HEPES, 30 NaHCO_3_, 10 MgSO_4_, 25 glucose, 5 Na-ascorbate, 3 Na-pyruvate and 2 Thiourea oxygenated with 95% O_2_ and 5% CO_2_, pH 7.3–7.4, and osmolality adjusted to 315–320 mOsm with sucrose. The brainstem tissue was blocked and embedded in 2% agarose in slicing aCSF solution. Coronal brain slices containing NAc in thickness of 300 μm were cut using a compresstome (VF-300, Precisionary Instruments LLC., MA, USA). The slices then were incubated with slicing aCSF at 32°C for 30 min, followed by another incubation session with holding aCSF containing (in mM): 92 NaCl, 2.5 KCl, 1.25 NaH_2_PO_4_, 2 CaCl_2_, 20 HEPES, 30 NaHCO_3_, 2 MgSO_4_, 25 glucose, 5 Na-ascorbate, 3 Na-pyruvate and 2 Thiourea oxygenated with 95% O_2_ and 5% CO_2_, pH 7.3–7.4, and osmolality adjusted to 310–315 mOsm with sucrose. The slices were transferred to a chamber mounted on an upright microscope equipped with Nomarski and epifluorescent optic systems (BX51WI, Olympus Optical Co., Ltd, Tokyo, Japan) and camera (Flash4.0 V3, Hamamatsu Photonics, Shizuoka, Japan). Then the slices were continuously perfused with regular aCSF that containing (in mM): 124 NaCl, 2.5 KCl, 1.25 NaH_2_PO_4_, 1 MgSO_4_, 2 CaCl_2_, 24 NaHCO_3_, and 11 glucose, oxygenated with 95% O_2_ and 5% CO_2_, pH 7.3–7.4, and with an osmolality of 305–310 mOsm. All of recordings were conducted at 30–32°C with aCSF perfusion at ∼2.5mL/min, and the electrodes were placed into NAc through the guidance of morphological landmarks and expression of viral fluorescent protein. Once the working MIE electrode was positioned in the NAc, a square wave was applied from 0 to 1 V at a frequency of 30 Hz (Incre. E = 0.004 V; Amplitude = 0.04 V) every 15 seconds for 30 min to prime, or habituate, the MIE to the brain slice environment. Next, the same square wave was applied for an additional 5 min and peak oxidative current was measured from each electrical potential scan as the baseline. Following the same baseline protocol, 10 Hz photostimulation, 10 ms pulse width was applied with a blue LED for 15 minutes while simultaneously applying the electrical potential square wave and peak oxidative current was measured. Finally, the LED was turned off and the square wave was applied for an additional 10 min.

### Statistics

All data are shown as the mean +/-standard error of the mean (SEM) unless otherwise noted. For comparing average calibration data (Figure 2D), a mixed effect model was used because all 10 electrodes were not tested at all six concentrations. A repeated measured ANOVA was used to compare the average peak oxidative current before and during bath application of dynorphin and to compare the average peak oxidative current before, during, and after 10 Hz photostimulation. Statistical analyses were performed in GraphPad Prism 9.

## Acknowledgment

This work was supported by the National Instituted of Health R00 DA038725 (RA), R21 DA048650 (RA), R01 NS123070 (RA), F32 DA053093-01A1 (SC), R01 NS117899 (JGM), NARSAD Young Investigator Grant from the Behavior Research Foundation, grant no. 28243 (RA). Special thanks to Huafang Li for her help with the scanning electron microscope.

## Notes

### Competing Interest Statement

The authors have declared no competing interest.

